# Optogenetic stimulation of Lbc GEF-mediated Rho activity dynamics promotes cell invasion

**DOI:** 10.1101/2025.03.28.646036

**Authors:** Jessica Wagner, Konstantina Feller, Nicole Schrenke, Nina Schulze, Annette Paschen, Leif Dehmelt, Perihan Nalbant

**Author notes:** **Contact:** Corresponding authors (P.N.); (L.D.), **Additional Footnote** Co-corresponding authors.

## Abstract

Cancer cell invasion relies on dynamic cell shape changes, which originate from protrusive and contractile intracellular forces. Previous studies revealed that contractile forces are controlled by positive-feedback amplification of the contraction regulator Rho by Lbc GEFs. These GEFs were previously linked to tumor progression, however, the underlying mechanisms are poorly understood. Here, we generated a mouse melanoma model, in which cytosolic levels of the Lbc GEF GEF-H1 are controlled by light. Using this model, we found that increased GEF-H1 levels strongly stimulate cell contraction dynamics. Interestingly, increased contraction dynamics rapidly induced expansion of tumor spheroids via a focal adhesion kinase-dependent mechanism. Furthermore, long-term stimulation led to the escape of individual cells from spheroids. These findings reveal new insights into the oncogenic roles of Lbc GEFs, and how they might promote tumor cell invasion. We propose a mechanism, in which increased cell contraction dynamics results in asymmetric pulling forces at the tumor border, promoting the detachment and escape of individual cells.

## Introduction

Directional cell migration is an essential process in the development and maintenance of multicellular organisms. In a simple model, cell migration can be described as a multi-step process: 1) protrusion of the leading edge, 2) formation of new adhesions near the leading edge, 3) increase of contractile forces and 4) disassembly of adhesions and cell retraction of the rear edge.^1^ These basic steps of protrusion, adhesion and retraction are controlled by dynamic signal networks that are modulated by numerous biochemical and mechanical inputs. The underlying signal networks have to control the coordination of cell protrusion, adhesion and retraction in space and time, and a loss of these control mechanisms can lead to dysregulated cell migration.

An important example of aberrant regulation of cell migration is cancer cell metastasis, which is a key step in the pathology of most cancer patients. During metastasis, cancer cells escape from the locally growing tumor and migrate into distant tissues or organs, where they then continue to proliferate. A critical step during metastasis is the local invasion of tumor cells into the adjacent tissues. This invasive cell movement is based on the formation of specialized actin structures, which can generate protrusive or contractile forces.^2,3,4^ A prominent role of cell contraction in this context was demonstrated by earlier studies that revealed significantly reduced tumor cell motility and invasiveness in several tumor cell models after inhibition of contraction signaling molecules.^5,6,7,8,9^

Cell contraction is mainly regulated by the small guanosine triphosphatase (GTPase) Rho. Previous studies revealed a mechanosensitive cell contraction signal network, in which Rho acts as the central mediator to generate subcellular myosin-based cell contraction pulses. In this signal network Rho activity is amplified by a fast-acting positive feedback loop, which is mediated by the Lbc-type guanine nucleotide exchange factor GEF-H1 and related Lbc-GEFs.^10^ Subsequently, Rho activity is inhibited by time-delayed slow negative feedback.^10^ The dynamics of the resulting contraction pulses are modulated by mechanical inputs from the cellular environment and by changes in the expression level of GEF-H1. ^10,11^ Interestingly, increased mRNA and protein levels of GEF-H1 were detected in aggressive cancers, including breast cancer and melanoma.^12,13,14,15,16,17^ Similar observations were made with other members of the Lbc family,^18,19^ which suggest that positive feedback amplification of Rho by Lbc-type GEFs is linked to metastatic behavior and advanced stage tumors. However, how these GEFs contribute to cancer progression is still poorly understood and requires further investigation, particularly as GEFs are also being considered as targets for cancer therapy.^20^

Here, we used an optogenetic strategy to establish a tumor spheroid model that enables direct, light-based control of the effective cytosolic concentration of the cell contraction regulator GEF-H1. This system does not only enable full temporal control of cell contraction at the level of entire spheroids, but also allows light-based control of cell autonomous, pulsatile contraction dynamics by modulating positive feedback amplification of Rho activity. Using this system, we found that multicellular spheroids expand during the optogenetic stimulation of cell contraction dynamics. Light-induced spheroid expansion required focal adhesion kinase (FAK) activity, suggesting that integrin-mediated cell-matrix interactions at the tumor periphery might play a role in this process. Furthermore, long-term stimulation of cell autonomous contraction dynamics also lead to the escape of individual cancer cells from the tumor spheroid. Based on these investigations, we propose that enhanced cell contraction dynamics in individual cells of the spheroid generate an outward-directed force at the interface between the spheroid periphery and the surrounding ECM, which promotes an invasive tumor cell phenotype.

## Results

### Development of a stable mouse melanoma cell system for optogenetic control of Rho activity amplification

Amplification of Rho activity by Lbc-type GEFs is a critical component of myosin-based subcellular contraction dynamics.^10^ To investigate the effects of increased contraction dynamics on tumor cell behavior, we established an optogenetic strategy in the B16F1 mouse melanoma cell model system. In our approach, we implemented the microtubule regulated Lbc-GEF GEF-H1.^21,22,23^ Similar to previous observations in U2OS cells,^10^ transient expression of the constitutively active, microtubule-binding deficient GEF-H1 mutant C53R, strongly stimulated myosin IIa based cell contraction pulses in B16F1 cells (Figures 1A-1D and Video S1). To enable accurate control of cell contraction dynamics, we combined the GEF-H1 C53R mutant with the optogenetic LOVTRAP system.^24,11^ The LOVTRAP system utilizes reversible, light-induced protein dissociation of the photo-sensitive LOV2 domain from *Avena sativa* phototropin 1 and a small protein called Zdark1 (Zdk1).^24^ We genetically fused the microtubule-binding deficient GEF-H1 C53R mutant to Zdk1 and expressed this construct with the mitochondria targeted LOV2 domain (NTOM20-LOV2).^11^ In cells, Zdk1 interacts with LOV2 in the dark state, sequestering GEF-H1 at the mitochondria and thereby depleting it from the cytosol. This interaction can be reversibly inhibited by blue-light (425-475 nm) illumination, which releases GEF-H1 into the cytosol and stimulates Rho activity amplification and actomyosin-based cell contraction pulses (Figure 1E).

**Figure 1:**
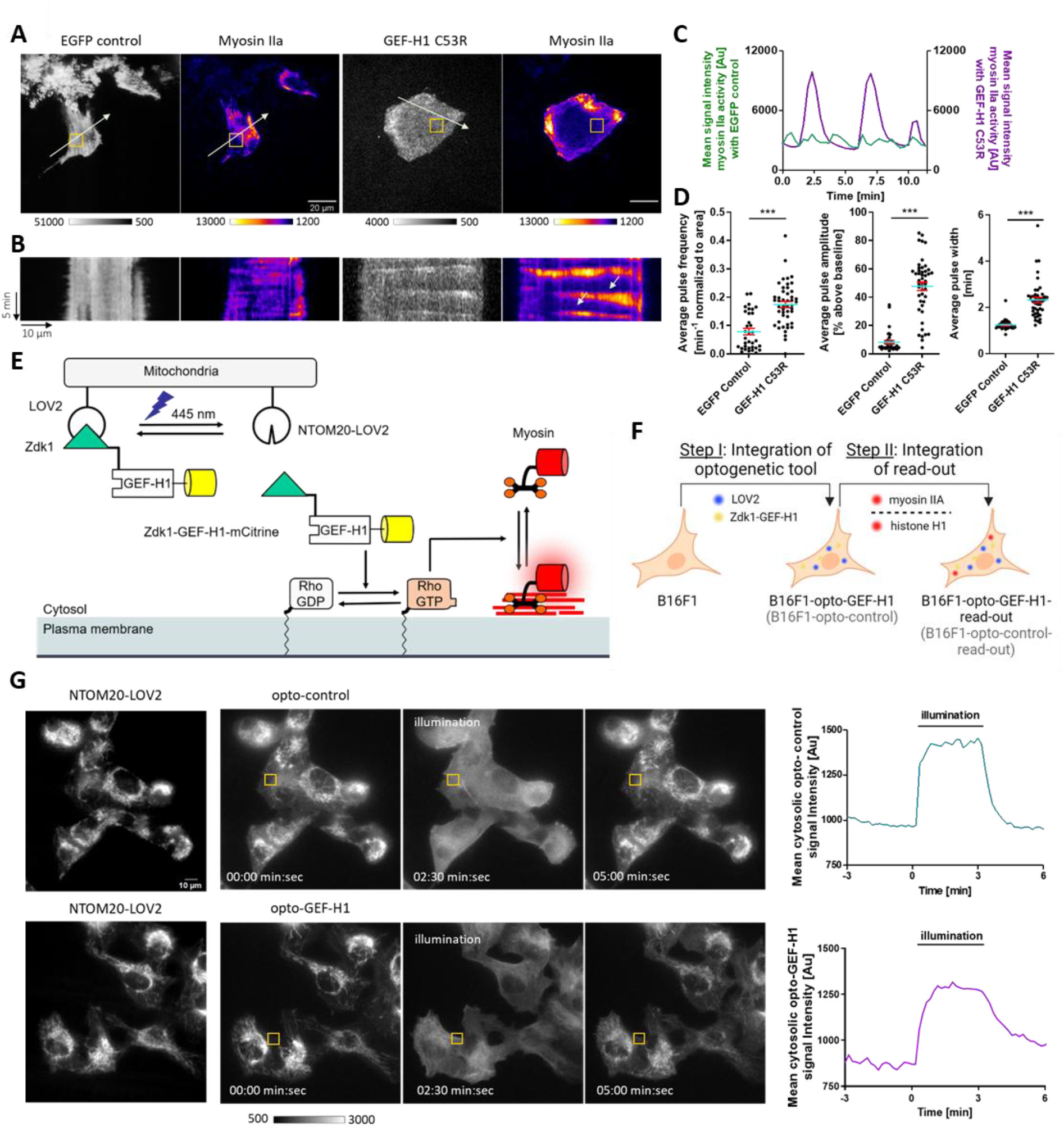
Development of a stable mouse melanoma cell system for optogenetic control of the cytosolic GEF-H1 concentration. (A-D) Increased expression of a constitutively active GEF-H1 mutant in B16F1 mouse melanoma cells leads to enhanced cell contraction dynamics. (A) Representative TIRF images of cells transiently expressing myosin IIa (pCMV-mCherry-MHC IIA) together with the constitutively active GEF-H1 C53R mutant (pCMV5-EGFP-GEF-HI C53R) or a control vector (pEGFP-N1), respectively. Scale bar = 20 μm. (B) Corresponding kymograph analysis along the arrows in A. (C) Myosin IIa signal intensity plots corresponding to the orange box in A. (D) Quantification of average contraction pulse frequency, amplitude and width. N=41 GEF-H1 C53R cells and N=24 control cells from 3 independent experiments, Error bars represent S.E.M., Unpaired t-test. (E) Schematic representation of optogenetic GEF-H1 release from the mitochondria into the cytosol and subsequent stimulation of cell contraction dynamics by myosin IIa. (F) Schematic representation of stepwise strategy to generate stable B16F1 cell lines expressing a GEF-H1-coupled optogenetic tool, the corresponding control and fluorescently tagged read-out proteins. (G) Left: Representative epifluorescence images showing the mitochondrial anchored photo-sensitive LOV2 domain (NTOM20-moxBFP-LOV2) and opto-control (mCitrine-Zdk1), or opto-GEF-H1 (mCitrine-Zdk1-GEF-H1 C53R) before, during and after optogenetic stimulation at 445-488 nm. Scale bar = 10 μm. Right: Intensity plots of opto-control and opto-GEF-H1 signal corresponding to the orange boxes in left panels.

To enable uniform light-based control of cell contraction dynamics in cell populations, we used the PiggyBac transposon system to generate monoclonal stable cell lines that express the mitochondrial-anchored LOV2 domain (NTOM20-LOV2) and the Zdk1-GEF-H1 C53R fusion protein or a control construct that lacks GEF-H1 (Figure 1F). We refer to these cell lines as B16F1-opto-GEF-H1 and B16F1-opto-control, respectively. Microscopy-based evaluation revealed efficient release of the control protein or the GEF-H1 fusion from mitochondria into the cytosol in a rapid and reversible manner upon blue light illumination (Figure 1G). We then stably integrated fluorescently tagged read-out proteins into these cell lines. In particular, we used myosin IIa to visualize cell contraction dynamics and histone H1 to track the cell position (Figure 1F).

### Dynamic cell contraction response to optogenetic modulation of Rho amplification in single cells

RhoA is one of the 3 best-studied, canonical Rho GTPases and is mainly responsible for controlling actomyosin based cell contraction. Upon activation by Lbc-type GEFs such as GEF-H1, Rho binds and activates the protein kinase ROCK, which in turn activates non-muscle myosin II isoforms.^25,26^ This results in contraction of actomyosin structures at the cell cortex and in the formation of stress fibers.^27^ Prior studies in human U2OS osteosarcoma cells, revealed prominent pulses of Rho activity and myosin contraction in the cell cortex upon elevation of cellular GEF-H1 levels.^10,28^

To study the effect of increased GEF-H1 levels on cell contraction dynamics in melanoma, we used the B16F1-opto-GEF-H1 and B16F1-opto-control cells. To investigate the effect of optogenetic GEF-H1 manipulations on cell contraction dynamics, we used total internal reflection fluorescence microscopy (TIRF-M) to measure plasma membrane recruitment dynamics of myosin IIa, which was also stably expressed as a fluorescently labeled fusion protein. We applied photoactivation sequences which consisted of three sequential steps: 1) Prerun: Myosin IIa imaging without blue light stimulation, 2) Perturbation: Myosin IIa imaging with blue light stimulation, resulting in optogenetically increased effective cytosolic GEF-H1 concentrations, 3) Recovery: Myosin IIa imaging without blue light stimulation.

In the first set of experiments, we optogenetically activated GEF-H1 for 5 min during the perturbation step. Shortly after starting this light-induced stimulation, we observed a rapid increase of myosin IIa signals at the plasma membrane (Figures 2A and 2B and Video S2). This indicates that GEF-H1 was able to stimulate Rho activity which in turn activated myosin, resulting in its increased association with cortical actin filaments near the plasma membrane. Even during prolonged blue-light illumination, the signal dropped sharply after an initial peak, presumably due to negative feedback regulation ^10^. The combination of rapid positive and slow negative feedback is known to stimulate oscillatory or excitable signal network dynamics in U2OS osteosarcoma cells.^10,28^ Here, we also observed the formation of single prominent myosin pulses and wave propagation through the cell, indicating that similar dynamics occur in B16F1 melanoma cells (Figures 2A-2C). Quantification of the relative change of myosin IIa signal intensity revealed a strong increase of myosin signal intensity when GEF-H1 was released from mitochondria (96.09±11.26%). Applying the same protocol to the opto-control cell line had no effect (Figure 2G).

**Figure 2:**
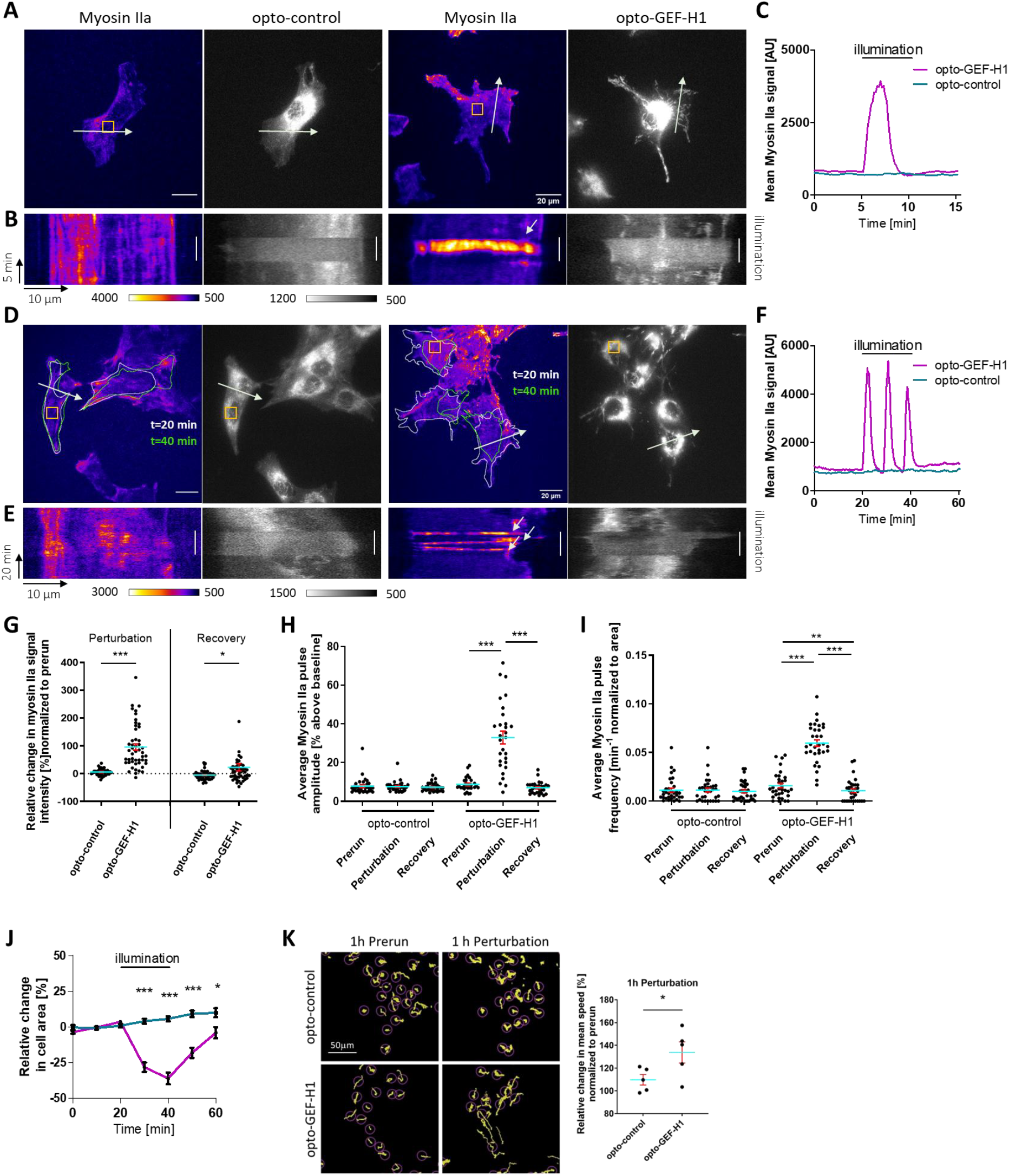
Direct optogenetic stimulation of pulsatory cell contraction dynamics in adherent mouse melanoma cells. (A, D) Representative TIRF images of myosin IIa signal and opto-control (left) or opto-GEF-H1 (right) with 5 min (A) or 20 min (D) optogenetic stimulation. Scale bar = 20 μm. (B, E) Corresponding kymograph analysis along the arrows in A and D. Optogenetic stimulation indicated by white line. (C, F) Intensity plots depicting myosin IIa signals in opto-control and opto-GEF-H1 cells, corresponding to the orange boxes in A and D. (G) Relative change in myosin IIa average signal intensity [%] during 5 min optogenetic perturbation and during recovery in the dark, normalized to prerun measurements before illumination. N≥44 cells per condition from 3 independent experiments, Unpaired t-test. (H-I) Quantification of contraction dynamics upon GEF-H1 release. Average amplitude (H) and frequency (I) of myosin-based contraction pulses without light (prerun), during optogenetic perturbation and during recovery. N ≥ 29 GEF-H1 cells and N ≥ 32 control cells from 3 independent experiments, Unpaired t-test. (J) Change in adherent cell area at individual timepoints relative to the interval before 20 min of optogenetic stimulation. Cell borders before optogenetic stimulation (t=20 min) and at the end of optogenetic stimulation (t=40 min) are indicated in (D) by white and green lines, respectively. N=42 cells and N=37 control cells from 4 independent experiments, Unpaired t-test. (K) Optogenetic stimulation of contraction dynamics leads to increased cell movement. (Left) Representative tracks (yellow) overlayed with nuclei detections at t=0min (magenta circles) of opto-control and opto-GEF-H1 cells, either during the prerun phase or during the optogenetic perturbation phase. (Right) Quantification of mean speed of opto-control and opto-GEF-H1 cells during 1h optogenetic perturbation relative to 1h prerun. N=5 experiments for each condition, Unpaired t-test. Error bars represent S.E.M.

Next, we extended the duration of optogenetically-induced GEF-H1 release to 20 min. This prolonged perturbation led to multiple prominent contraction pulses in B16F1-opto-GEF-H1 cells (Figures 2D-2F and Video S3). Both, the amplitude and frequency were significantly increased compared to the unperturbed state (Figures 2H and 2I.). These pulsatory dynamics were cell autonomous and differed in frequency and phase between neighboring cells. Myosin intensity dynamics returned to prerun levels shortly after starting the recovery phase, in which GEF-H1 was recaptured at mitochondria in the absence of blue light illumination. Together, these observations emphasize that an increase of cytosolic GEF-H1 levels can rapidly stimulate the cell contraction signal network dynamics.

In addition to the increase of contraction dynamics during optogenetic manipulation, we noticed that cells often started to round up. To quantify this observation, we determined the adherent area of individual cells throughout the three distinctive imaging phases (Figure 2J). Compared to opto-control cells, the average adherent cell area was significantly decreased by 35.93±4.07% after 20 min of optogenetic GEF-H1 release (Figure 2J). During the subsequent recovery phase, this effect was largely reversed as cells quickly began to reattach. Using the live-cell nuclear marker (Spy650-DNA, Spirochrome) and epi-fluorescence imaging, we also found that average cell movements significantly increased upon light-induced GEF-H1 release (Figure 2K). Taken together, these results show light-inducible and reversible optogenetic modulation of cytosolic GEF-H1 levels associated with increased cell contraction dynamics and cell movements in the melanoma model system.

### Response of melanoma spheroids to optogenetic manipulation of cell contraction dynamics

Next, we investigated how alterations of cell contraction dynamics on the level of individual cells might affect tumor related cell behavior in 3D. To address this question, we utilized well-established protocols to generate multicellular tumor spheroids (MCTS) that were embedded in a collagen matrix. ^29^ This system resembles the natural tumor environment more closely by including cell-cell and cell-matrix interactions. To study the effect of light-induced stimulation of cell contraction dynamics, we applied similar photoactivation protocols as those described above.

First, we used spinning disk confocal microscopy (SDCM) to investigate the overall cellular organization within spheroids of B16F1-opto cells that stably express fluorescently labelled histone H1. This enabled us to visualize cell nuclei within the spheroid and confirmed a uniform, densely packed architecture throughout the entire spheroid (Figure 3A). During the application of a short (5 min) optogenetic stimulation, we observed enhanced cell movement dynamics within the B16F1-opto-GEF-H1 spheroids (Video S4). To quantify this, we tracked the labelled nuclei over time and calculated the average movement velocity as a function of time. In the absence of blue light illumination only weak, baseline cellular movements were detected. However, in response to the optogenetic release of GEF-H1 from mitochondria in B16F1-opto-GEF-H1 spheroids, the average cell velocity immediately increased. In contrast, we did not observe such increase of dynamics in control spheroids (B16F1-opto-control) (Figure 3B).

**Figure 3:**
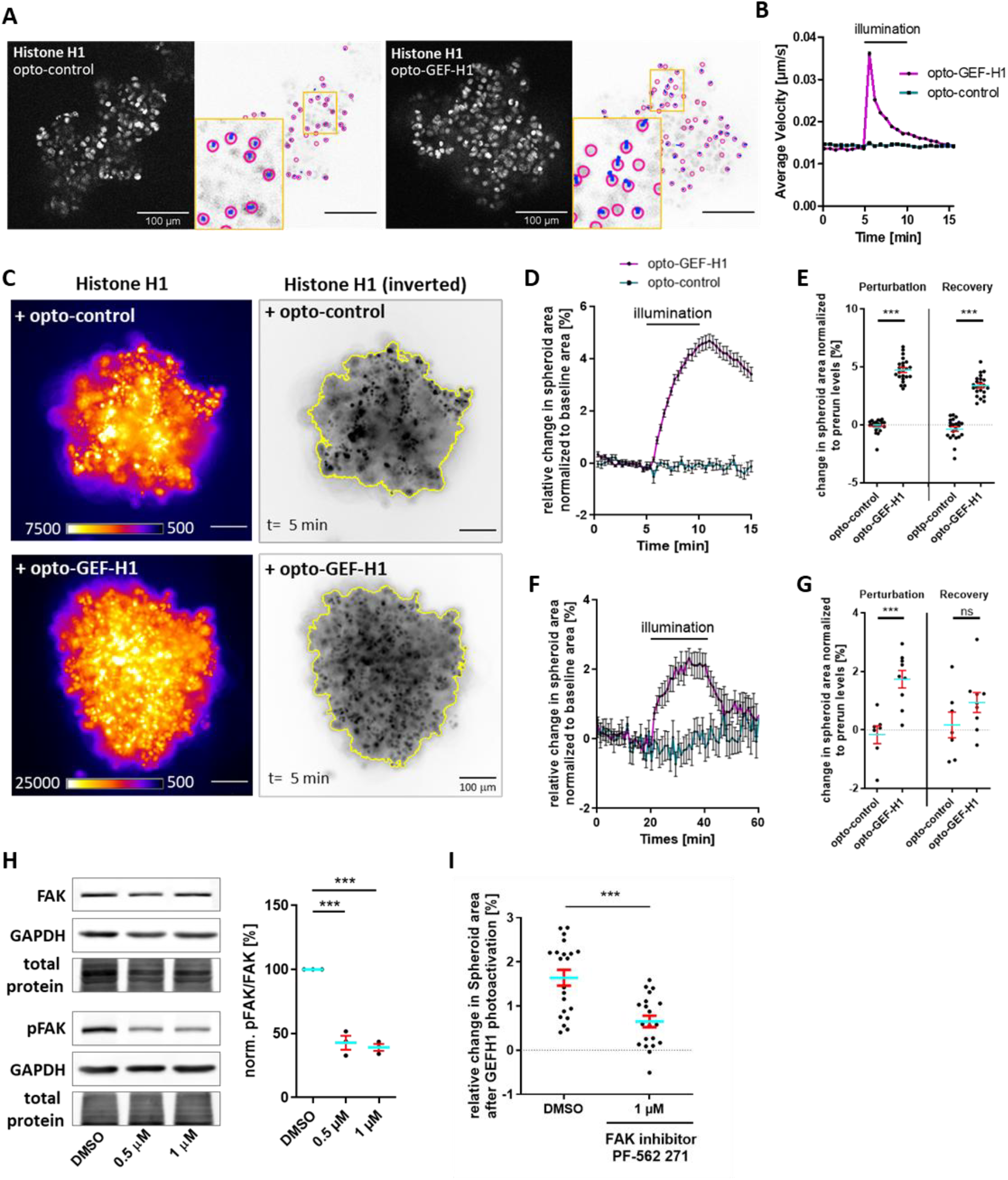
Optogenetic stimulation of contraction dynamics leads to expansion of multicellular melanoma spheroids. (A) Representative spinning disk confocal microscopy images of nuclei (Histone H1) in B16F1-opto-control or B16F1-opto-GEF-H1 spheroids. Neighboring panels show corresponding tracking paths (blue lines) of nuclei highlighted in magenta. Scale bar=100 μm. (B) Average movement velocity of opto-control (turquoise) and opto-GEF-H1 (magenta) cells within spheroids before, during and after illumination between timepoints t=5 and t=10 min. N=1978 cell tracks in 26 opto-GEF-H1 spheroids and N=1822 cell tracks in 23 control spheroids from 3 independent experiments. (C) Left panels: Representative epifluorescence images showing nuclei in opto-control (upper panels) and opto-GEF-H1 (bottom panels) spheroids. Right panels: Corresponding spheroid areas (yellow lines) before (t=5 min) optogenetic stimulation. Scale bar=100 μm. (D-G) Quantification of relative change in spheroid area upon optogenetic stimulation of contraction dynamics. (D, F) Relative change in spheroid area over time. Optogenetic stimulation (D) from t=5 min to 10 min and (F) from t=20 min to 40 min. (E, G) Relative change in spheroid area after 5 min optogenetic stimulation and after 5 min recovery (E) or after 20 min optogenetic stimulation and after 20 min recovery (G) normalized to prerun levels. (D-E): N=23 opto-GEF-H1 spheroids, N=21 opto-control spheroids from 3 independent experiments, Unpaired t-test. (F-G): N=9 opto-GEF-H1 spheroids, N=8 opto-control spheroids from 3 independent experiments, Unpaired t-test. Error bars represent S.E.M. (H-I) Spheroid expansion upon optogenetic stimulation of contraction dynamics requires FAK activity. (H) Left: Representative Western blot depicting FAK inhibitor PF-562 271-mediated inhibition of FAK autophosphorylation in B16F1 cells. Right: Quantification of FAK autophosphorylation relative to control DMSO treatment. n = 3 independent experiments. (I) Plot showing the relative change in spheroid area after 5 min optogenetic stimulation of spheroids treated with 1 μM of FAK inhibitor PF-562 271 or DMSO as vehicle control. DMSO: N=21 spheroids, FAK inhibitor: N=20 spheroids from 3 independent experiments. Unpaired t-test.

Next, we investigated how increased contraction dynamics and movement on the single cell level might affect the overall structure and dynamics of spheroids. Upon optogenetic GEF-H1 release (Figures 3C-3E and Video S5) in B16F1-opto-GEF-H1 spheroids, we observed a significant increase of the projected spheroid area while no change was measured in control spheroids (Figures 3C-3E). During the recovery phase, the projected area of GEF-H1 spheroids decreased again. To evaluate the spheroid expansion and regression kinetics in more detail, we performed prolonged imaging experiments and extended the prerun, optogenetic GEF-H1 stimulation and recovery times to 20 min. We found that spheroid area expanded over the entire illumination time (Figures 3F and 3G), and it was only during the subsequent 20 min recovery phase that the projected area decreased again nearly to control levels. Furthermore, repeated 5 min illumination pulses over the course of several hours or a single, 3h long illumination pulse induced fully reversible spheroid expansions that closely correlated with the illumination phases (Figure S1). Taken together, our experiments demonstrate the full optogenetic control that is enabled by our system, which allows both highly synchronous contraction pulses in cell populations using the short, 5 min pulses (Figures 2A-2C, Figures 3C-3E and Figures S1A, B), as well as cell autonomous, asynchronous pulses using longer pulse durations (Figures 2D-2F, Figures 3F and 3G and Figures S1C, D). Furthermore, our experiments clearly show that spheroid expansion depends on continuous stimulation of cell contraction dynamics. Interestingly, both the increase (t_1/2_ ≈ 2.5 min) and the decrease (t_1/2_ ≈ 10 min) of the projected spheroid area were substantially slower compared to the near instantaneous increase of single cell movements. This suggests that spheroid expansion is not a direct consequence of increased single cell movements, but rather occurs subsequently in a downstream process or via a different process that acts in parallel.

The expansion of the multicellular spheroids upon stimulation of cell contraction in 3D was initially surprising, as analogous experiments with single cells on a 2D surface showed a reduction of the cell adhesion area (Figure 2). However, the 2D and 3D experiments differ quite significantly, in particular concerning interactions between cells and their environment (Figure S3). In 2D, cells adhere to the surface in a relatively isotropic fashion and induction of cell contraction leads to a similarly isotropic contraction and reduction of the cell adherent area. In 3D, cells in the spheroid center are also expected to form relatively isotropic adhesions, however, in this case these adhesions mainly occur between cells and not with the extracellular matrix. In contrast, cells at the spheroid border are expected to form highly anisotropic interactions – either with other cells towards the spheroid center, or with the surrounding matrix towards the spheroid periphery. We hypothesized that the observed spheroid expansion upon GEF-H1 release could result from these anisotropic interactions: A stronger interaction of border cells with the surrounding matrix compared to other cells towards the center would lead to an imbalance of force transduction during cell contraction. This imbalance could lead to a net pulling force at the spheroid periphery, which could drive spheroid expansion (Figure S3).

If this hypothesis is correct, an inhibition of the interaction between the border cells and the surrounding matrix should reduce spheroid expansion upon stimulation of cell contraction. As cell-matrix interactions are highly dependent on focal adhesions (FA), we hypothesized that spheroid expansion should be dependent on these structures. FAs mediate cell-matrix interactions by a mechanism that involves the activation of the non-receptor tyrosine kinase FAK (focal adhesion kinase) downstream of transmembrane integrin receptors. To test this hypothesis, we inhibited FAK using the small molecule compound PF-562 271 (Figure 3H). Similar to experiments without drug treatment (Figures 3C-3E), control spheroids pretreated with DMSO expanded upon optogenetic stimulation (Figure 3I). In contrast, enlargement was significantly reduced in spheroids which were treated with the FAK inhibitor (Figure 3I), showing that spheroid expansion indeed requires FAK-dependent cell substate adhesions.

Cell adhesion also plays an important role in cell migration. However, to enable efficient migration, cells have to control their adhesion very dynamically. Previous studies found that a constant increase in cell contraction leads to more stable adhesion, which would not be compatible with efficient cell migration. The activity of Lbc-type GEFs like GEF-H1 on the other hand stimulates highly dynamic cell contractions. To assess the effects of increased GEF-H1 activity on focal adhesion dynamics we transiently transfected individual adherent cells with fluorescently tagged paxillin and measured the average FA lifetime. We found that FA lifetimes in B16F1-opto cells were significantly reduced during light-triggered GEF-H1 activity stimulation compared to the initial, prerun stage (Figures S2A and S2B), showing that active GEF-H1 can increase focal adhesion turnover.

To escape from the primary tumor, cells have to increase their migratory and invasive capacity. Both cell migration and invasive behavior require myosin-based cell contractions and dynamic cell-ECM interactions.^30,31^ Interestingly, the results above suggest that Lbc-type GEFs can stimulate both of these dynamic processes. Furthermore, the expression of multiple Lbc-type GEFs is increased in advanced tumors, including GEF-H1, LARG and p190 Rho GEF.^12,14,16,18,19^ We therefore wondered how a longer-term increase of Lbc-GEF-induced contraction dynamics might affect the overall behavior of tumor spheroids. To address this, we used the established light-controlled melanoma spheroid system and extended the duration of the illumination over several hours. Strikingly, we frequently observed the escape of individual tumor cells during such long-term optogenetic stimulations of B16F1-opto-GEF-H1 spheroids (Figures 4A-4C and Video S6). The escape events occurred throughout the 12 h of optogenetic stimulation, starting already after 2 h (Figure 4C). In contrast, segregation of cells was very rarely observed in the control conditions (Figure 4C). Taken together, these results show that enhanced cell contraction dynamics stimulated by Lbc-type GEF-mediated Rho activity amplification can promote cancer cell detachment from tumor spheroids and increase the invasive behavior of mouse melanoma cells.

**Figure 4:**
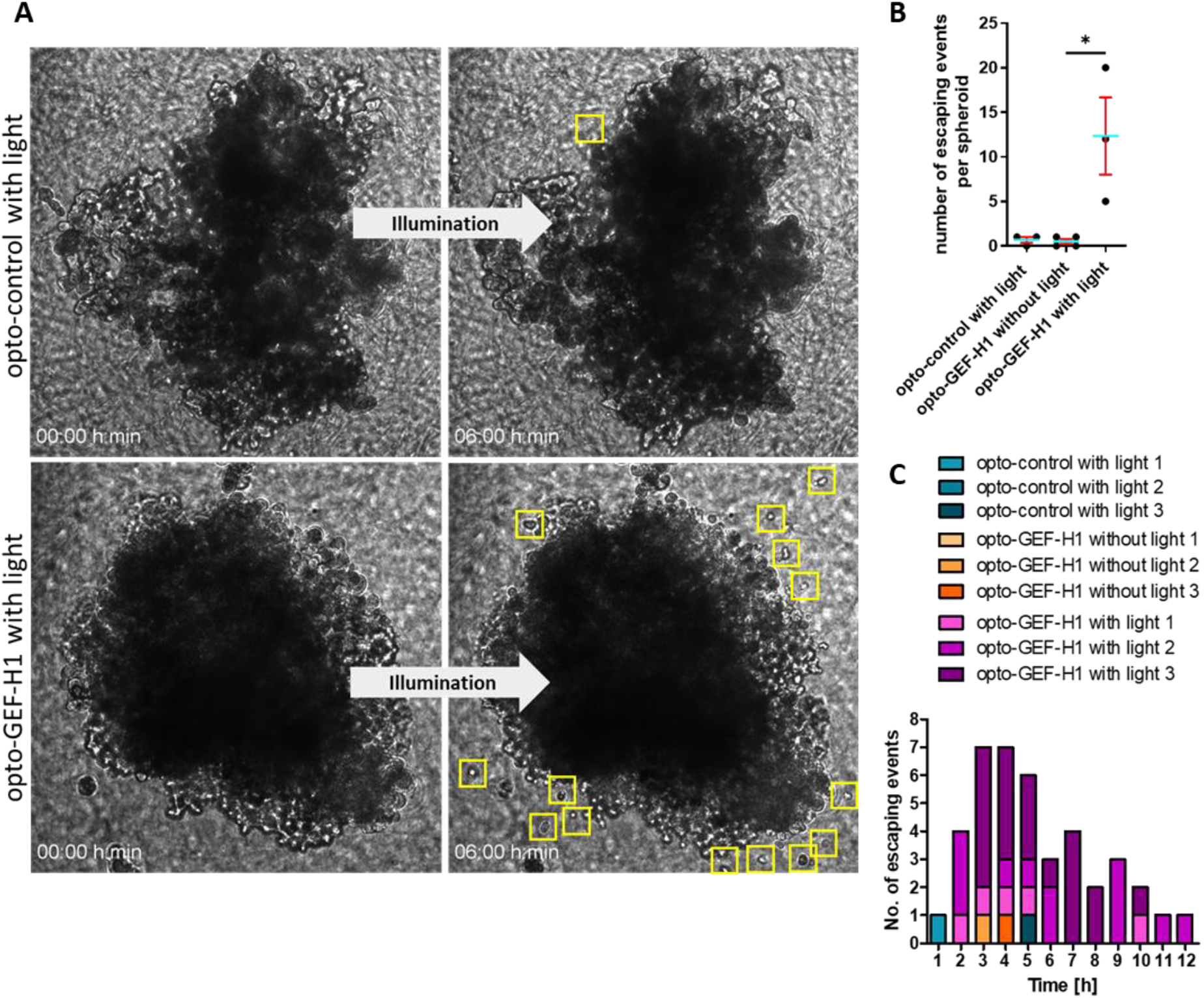
Optogenetic stimulation of contraction dynamics leads to detachment of individual cells from melanoma spheroids. (A) Representative bright field microscopy images of B16F1-opto-control (upper panel) and B16F1-opto-GEF-H1 (lower panel) spheroids embedded in a collagen matrix before (left) and after (right) continuous optogenetic stimulation. Cells that detached from the spheroids are marked by yellow boxes. Scale bar=100 µm. (B) Total number of cell-escaping events from B16F1-opto-GEF-H1 spheroid with optogenetic stimulation and control conditions either without illumination or using B16F1-opto-control spheroids. (C) Chronological sequence of escaping events measured for 3-4 spheroids for each condition from 3 independent experiments.

## Discussion

In this study, we investigated the role of cell contraction signal network dynamics in tumor cell behavior by establishing a monoclonal light-controlled mouse melanoma model. With this model we were able to directly control Rho activity amplification and myosin-based contraction dynamics with high temporal resolution in 2D cell culture and multicellular 3D tumor spheroids. Our studies revealed new insights into mechanisms, how increased cell contraction dynamics in tumors could promote invasive and metastatic cancer progression.

Multicellular tumor spheroids (MCTS) recapitulate key aspects of the multi-layered 3D architecture of tumors. In contrast to planar 2D cultures, cells within spheroids are exposed to different environments depending on their location. At the most basic level, at least two distinct cell populations exist within the spheroid: cells within the core that mainly interact with each other via cell-cell contacts, and cells in the periphery that can also interact with the surrounding matrix (Figure S3). We hypothesized that this heterogeneity might play an important role in the expansion of spheroids that we observe after stimulation of cell contraction dynamics. In particular, cells at the spheroid border can generate two distinct contacts: cell-cell contacts towards the spheroid structure and cell-matrix contacts between the spheroid and the surrounding collagen polymers (Figure S3). This heterogeneity of contacts could lead to an asymmetry in force transduction between these regions of the spheroid, creating a directional force that pulls the border cells towards the surrounding matrix. In agreement with this idea, we observed a significantly reduced expansion after pharmacological inhibition of the focal adhesion regulator FAK. FAK was previously found to be involved in tumor related cell behavior in multiple ways, for example by stimulating the dynamic turnover of focal adhesions.^32^ The counterforce to cell-ECM adhesions that preserves spheroid integrity is generated by Cadherin-mediated cell-cell contacts. In B16F1 cells, this cell-cell contact based counterforce is expected to be relatively weak, as E-Cadherin levels were comparably low in this cell type.^33^ This imbalance between weak intercellular forces that hold the spheroid together and stronger outward directed forces mediated by cell-matrix contacts might facilitate spheroid expansion and ultimately lead to separation of individual cells from the tumor mass (Figure S3).

Optogenetic manipulation of GEF-H1 levels in our light-controlled melanoma model system did not only increase basal-level contraction, but very prominently stimulated pulsatile contraction dynamics (Figure 2D-2F; 2H and 2I). Our earlier studies showed that these dynamics are due to a combination of fast positive and slow negative feedback regulation.^10^ Increased contraction dynamics could have a profound effect on the function of focal adhesions. In general, FA can adopt two different states in response to myosin-based contraction and matrix stiffness: an undynamic, stable or a dynamic, tugging state.^34^ The highly pulsatory character of contraction dynamics induced by Lbc-type GEF might notably stimulate FA dynamics by repeatedly applying local tugging forces, which could increase force transduction.^35,36^ In addition, dynamic changes in the forces applied to FAs might enable cells to locally probe the physical properties of their environment and to find optimal migration routes. Via such a mechanism, increased contraction pulse frequency and amplitude could be a critical component for efficient invasive cell movements.^35,36^

Our current studies focused on GEF-H1, however, this molecule might not be unique in its ability to stimulate such a pro-invasive phenotype. Conceptually, all members of the Lbc-type GEF family might be able to stimulate pulsatory cell contraction dynamics, as they all can mediate positive feedback amplification of Rho activity. In previous studies, stimulation of Rho activity dynamics was indeed shown for the Lbc-type GEFs GEF-H1 and LARG (Leukemia-associated Rho guanine nucleotide exchange factor; ARHGEF12).^10,28^ For LARG in particular, overexpression was shown to transform mouse fibroblasts.^37^ and elevated expression levels were found in invasive edges of glioblastoma tissue.^18^ Furthermore, the Lbc-type GEF p115RhoGEF (ARHGEF1) was found to promote cell migration and amoeboid-like morphology of MMP inhibitor treated glioblastoma.^38^ Thus, increased Lbc-GEF signaling can be linked to the progression of several types of invasive tumors and this link might be related to increased cell contraction dynamics.

Cancer cells are often characterized by their ability to dynamically switch between different migration modes in order to achieve efficient movement through their environment. In our study, the invasive melanoma cells that detach from the tumor spheroid, displayed a relatively compact morphology. This morphology in compatible with a fast, matrix degradation independent amoeboid-like cell migration mode.^39^ Indeed, GEF-H1 and other Lbc-type GEFs were previously reported to promote amoeboid-like cell migration.^40^ Amoeboid-like migration strongly relies on myosin-based cell contraction that enables cells to squeeze themselves through matrix pores, enabling fast migration in 3D.^41,42,43^ Interestingly, in this migration mode, myosin-based contractions are not only fundamental for the retraction of the cell rear, but also for protrusion of the cell front via the dynamic generation of membrane blebs.^42^ Our observations therefore suggest that the GEF-H1-induced enhancement of contraction dynamics promotes amoeboid-like migration in our 3D melanoma model to enable fast and matrix degradation independent invasion.

In conclusion, we generated a light-controlled mouse melanoma model to rapidly stimulate cell contraction dynamics by modulating the effective cytosolic GEF-H1 concentration. We show that enhanced contraction dynamics lead to cell rounding and increased focal adhesion turnover. Within a 3D collagen matrix, induction of contraction dynamics results in global expansion of multicellular tumor spheroids and scattering of individual cells. We propose that this escape results from the unique properties of Lbc-type GEFs, which amplify Rho activity, stimulate pulsatory contraction dynamics and cell-matrix contact turnover, leading to asymmetric forces at the tumor border that enable separation of individual cells from the tumor mass.

## Materials and methods

### Plasmids

The expression vectors for the GEF-H1 C53R mutant (pCMV5-EGFP-GEF-H1-C53R),^44^ the fluorescently tagged myosin IIa (pCMV-mCherry-MHC IIA;^45^ Addgene plasmid #35687) and control EGFP (pEGFP-N1; Clontech Laboratories) were described previously.

The PiggyBac constructs used to generate stable cell lines expressing the optogenetic tool were constructed starting from vectors established previously.^46^ To generate the PiggyBac vector pPBCAG-NTOM20-moxBFP-GS-LOV2-IRES-Puro for expression of the mitochondrial-targeted LOV2 domain, the fluorescent moxBFP tag (Addgene plasmid #68064;^47^) was amplified together with a GSGSGS linker sequence via PCR and inserted into pTriEx-NTOM20-mVenus-LOV2 (kind gift from Klaus Hahn, University of North Carolina, Chapel Hill, USA)^24^ using BamHI and BseRI restriction sites and Gibson assembly. In a second step, the NTOM20-moxBFP-GS-LOV2 sequence was amplified via PCR and inserted into pPBCAG-cHA-IRES-Puro (kind gift from Christian Schröter, Max-Planck-Institute Dortmund, Germany;^46^) via the NotI and XhoI restriction sites and Gibson assembly.

To generate the PiggyBac perturbation vectors pPBCAG-mCitrine-Zdk1-IRES-Neo (control) and pPBCAG-mCitrine-Zdk1-GEF-H1-C53R-IRES-Neo, intermediate constructs were first generated starting from the previously published pTriEx-mCherry-Zdk1-GEF-H1 C53R vector.^28^ First, human GEF-H1 C53R was removed using the EcoRI and HindIII restriction sites, and replaced by mouse GEF-H1 (wt) via Gibson assembly, after amplification from pCMV-SPORT6-GEF-H1 (Horizon discovery/Dharmacon #MMM1013-202762494). Next, mCherry was replaced with mCitrine (mCitrine-C1; Addgene plasmid #54587) using the AgeI and HindIII restriction sites and Gibson assembly. The C53R point mutation was introduced by PCR-mutagenesis (primers: 5’-CGGATCGAATCCCTCACTC-3’; 5’-CTCTGGTGGTCGTCTTACTTCG-3’; Eurofins). Finally, mCitrine-Zdk1 or mCitrine-Zdk1-GEF-H1 C53R (mouse) were amplified via PCR and inserted into pPBCAG-rtTAM2-IN (kind gift from Christian Schröter; Max-Planck-Institute Dortmund, Germany) (Addgene plasmid #124166;^48^) using Gibson assembly.

To generate the PiggyBac readout vector pPBCAG-mCherry-MyosinIIA-C18-IRES-Hygro, mVenus in mVenus-MyosinIIA-C-18 was first replaced with mCherry from mCherry-NMHCIIA (Addgene plasmid #35687)^45^ via Gibson assembly using the HindIII and KpnI restriction sites. In parallel, the puromycin resistance in pPBCAG-cHA-IRES-Puro was replaced by the hygromycin resistance gene (from pBabe-Hygro; Addgene plasmid #1765)^49^ via PCR amplification and Gibson assembly using the ClaI and MscI restriction sites. Finally, mCherry-MyosinIIA-C-18 was inserted into pPBCAG-cHA-IRES-Hygro using NotI and XhoI sites via Gibson assembly. The PiggyBac readout vectors pPBCAG-mCherry-HistoneH1-IRES-Hygro and pPBCAG-mCherry-CaaX-IRES-Hygro were generated similarly via Gibson assembly by inserting mCherry-Histone1 mouse or the plasma membrane targeting sequence of K-Ras mCherry-CaaX, respectively, into pPBCAG-cHA-IRES-Hygro using the NotI and XhoI restriction sites.

### Cell Culture and reagents

B16F1 mouse melanoma cells (Kind gift from Annette Paschen, University Hospital Essen, Germany) were cultured in RPMI 1640 medium (Life technologies; Gibco) supplemented with 10% fetal bovine serum (Life technologies; Gibco) at 37°C and in 5% CO_2_ humidified atmosphere. During cell splitting, 0.05% Trypsin/EDTA (Pan Biotech) was used for trypsinization.

To generate stable cell lines, the PiggyBac transposon system was utilized.^46^ To generate B16F1-opto-GEF-H1, B16F1 cells were transfected with three plasmids: pPBCAG-NTOM20-moxBFP-GS-LOV2-IRES-Puro, pPBCAG-mCitrine-Zdk1-GEF-H1-C53R-IRES-Neo and the PB transposon vector pBase (Kind gift from Christian Schröder, Max-Planck-Institute Dortmund).^46^ To generate the control cell line (B16F1-opto-control), pPBCAG-mCitrine-Zdk1-IRES-Neo was used instead of pPBCAG-mCitrine-Zdk1-GEF-H1-C53R-IRES-Neo. Selection was performed with 1.5 µg/ml Puromycin (InvivoGen) and 1250 µg/ml G418 (Geneticin; InvivoGen). After 14 days of selection, monoclonal cell colonies were obtained via fluorescence activated cell sorting (FACS). The established B16F1-opto-GEF-H1 and B16F1-opto-control cell lines were subsequently transfected with pPBCAG-mCherry-MyosinIIA-C18-IRES-Hygro or pPBCAG-mCherry-HistoneH1-IRES-Hygro, respectively and selected with 125 µg/ml Hygromycin B Gold (InvivoGen) for 14 days, before a second round of FACS analysis. For culturing, the cell culture medium was continuously supplemented with the corresponding antibiotics at these concentrations.

For spheroid generation, B16F1 cells (3000 cells in 100 μl growth medium/pro well) were seeded on 96-well cell culture plates (Sarstedt) coated with 1% agar in PBS (Agar, Noble Agar, Ultrapure; Thermo Scientific). After 72 h, B16F1 spheroids were harvested and integrated into a collagen matrix essentially as described previously ^29^ (protocol kindly provided by Alexander Roesch, University Hospital Essen).

### Transfection and pharmacological treatments

Transient and stable transfection of plasmid DNA was performed using Lipofectamine 2000 according to the suppliers’ protocol (Life Technologies; Gibco). Pharmacological treatments with the FAK inhibitor PF-562 271 (0.5 µM, 1 µM; Sigma-Aldrich, PZ0387) were performed 6 h after transfection and at least 16 h before imaging. Pharmacological treatments of spheroids with the FAK inhibitor PF-562 271 (1 µM; Sigma-Aldrich, PZ0387) were performed 2 h after integration and at least 16 h before imaging.

### Microscopy and optogenetic manipulation of cytosolic GEF-H1 levels

For live-cell microscopy, cells were seeded on glass bottom dishes (MatTek) functionalized with 0.001% (v/v) collagen type I (collagen from calf skin, 0.1% solution in 0.1 M acetic acid; Sigma-Aldrich) in DPBS for 1 h at 37°C. For imaging at ambient CO_2_ levels, the culturing medium was replaced with imaging medium (RPMI 1640 (no phenol red); 10 % (v/v) FCS; 10 mM HEPES; 1 mM CaCl_2_; 1 mM MgCl_2_). Live-cell epifluorescence and total internal reflection fluorescence (TIRF) microscopy was performed on the Nikon Ti2-E TIRF DualCam system, equipped with H-TIRF for fully automated TIRF adjustments, an Andor Technology 2X iXon Life DU-888 back-illuminated EMCCD camera, the diode laser cw LuXX 488 nm (250mW), the DPSS laser cw Jive 561nm (300mW) and the Omicron Light Hub-4 compact laser beam combiner with AOTF laser modulator. The following emission filters and dichroic mirrors were used: Emission Filter#1: EGFP, 525/50 nm; Emission Filter #2: RFP, 600/50nm; Emission filter #3: Cy5, 705/72nm, NSTORM QUAD dicroic mirror (Reflection at 497-553 nm and 575-628 nm, Transmission at 523.5/42 nm; and 603/44 nm). Single cells were imaged using a CFI Apochromat TIRF 60x/1.49 oil immersion objective. Spheroids and single cells for 2D tracking experiments were imaged using a CFI Plan Apochromat λ 20x/0.75 NA air objective using the Nikon Imaging Software NIS elements Advanced Research. Confocal spinning disk microscopy of spheroids was performed using an Eclipse Ti-E (Nikon) inverted microscope with an Andor AOTF Laser Combiner, a CSU-X1 Yokogawa spinning disk unit and an iXon3 897 single photon detection EMCCD camera (Andor Technology). The following laser lines were used for excitation: EGFP: 488 nm (50 mW); RFP: 561 nm (50 mW). Images were acquired using a CFI APO TIRF 100×/1.49NA oil immersion objective (Nikon). Acquisition was controlled by Andor IQ Software (Andor Technology).

Optogenetic manipulation of the cytosolic GEF-H1 concentration via the LOVTRAP system was achieved by illumination with blue light (HC LED 465 filter with 460/60 nm excitation wavelength range). To avoid phototoxicity, we determined the minimal amount of blue light necessary for optogenetic GEF-H1 release. To achieve this, we used a 1000x neutral density filter in the 460/60 nm excitation light path. To combine image acquisition and photoactivation, the sample was exposed to blue light only between camera exposures. To avoid optogenetic release during brightfield imaging, a long-pass blocking filter (OG 550) was installed in the transmitted light path.

### Western blot analysis

Cells were washed once with ice-cold PBS and lysed in ice-cold RIPA buffer (50 mM Tris, pH 7.5, 150 mM NaCl, 1% NP-40, 0.25% sodium deoxycholate, 1 mM EDTA, 1× protease inhibitor cocktail, 1×phosphatase inhibitor cocktail, 1 mM Na3VO4 and 1 mM NaF). Cell debris were removed by centrifugation at 14,000 g for 20 min at 4°C. Protein concentration was determined using the Bradford assay. Equal amounts of total protein were mixed with 5× Laemmli sample buffer, boiled at 95°C for 10 min and separated by SDS-PAGE. After electrophoresis, proteins were transferred onto a PVDF membrane using a semidry blotter. Blots were blocked for 60 min at RT with Intercept® (TBS) Blocking Buffer (1:2, LI-COR Biosciences) and incubated overnight at 4°C with a primary antibody that recognizes FAK (1:1000 dilution; 05-537, clone 4.47; Millipore) or pFAK (1:1000 dilution; 44624G; Invitrogen) in blocking solution. Membranes were washed three times with TBS-T and incubated with the IRDye 800CW Goat anti–mouse secondary antibody (LI-COR Biosciences, 1:20000 dilution) for FAK detection and IRDye 800CW Goat anti–rabbit secondary antibody (LI-COR Biosciences, 1:20000 dilution) for pFAK detection for 1 h at RT. After additional washing steps with TBS-T and TBS (20 mM Tris, pH 7.6, and 137 mM NaCl) detection was carried out using the Odyssey scanner at 680 nm and 800 nm (LI-COR Biosciences).

### Image processing and data analysis

Microscopy images were processed using ImageJ/Fiji Software.^50^ The image stabilizer plugin (K. Li, “The image stabilizer plugin for ImageJ,” http://www.cs.cmu.edu/~kangli/code/Image_Stabilizer.html, February, 2008) was used to correct lateral drift. Representative microscopy images were adjusted with respect to brightness levels, cropping, scaling and false color-coding via look-up tables. All images that correspond to conditions that were compared against each other were treated identically.

### Cell contraction signal network dynamics analysis

To analyze myosin-based contraction dynamics, average pulse frequency, amplitude and duration were determined using an automated analysis tool that was described previously.^10^ Briefly, time-lapse movies with a frame rate of 3 per min were analyzed, in which individual cells were isolated and masked. The image resolution was reduced by a factor of 15-20 with averaging to reduce noise. To exclude signal changes that originate from dynamic cell protrusions, peripheral pixels surrounding individual cells were removed via a binary erode filter with a neighborhood count of 1. Global intensity changes were eliminated by division of each frame with the average background-corrected intensity. The resulting images were evaluated regarding their temporal intensity changes, by calculating several properties. 1) To measure peak amplitude, intensity maxima and minima in individual pixels were determined over time and the difference between a maximum and the preceding and following minima was calculated. To reduce the impact of noise, only amplitudes ≥ 1% were considered for the determination of the average peak amplitude. 2) To measure the pulse frequency and pulse width, only strong peaks with an amplitude above 10% (Figure 1D) or 20% (Figure 2I) were included. Pulse width was defined as the time difference between the preceding and following minima.

### Global contraction analysis

To measure changes in cell adhesion area, each cell was outlined using the wand (tracing) tool of ImageJ/Fiji ^50^ at given time points (t=0, 10, 20, 30, 40, 50, 60 min). Cells were only included for analysis if the entire area was detected in the imaging field throughout all time points. The measured area was normalized to the average area of the corresponding cells in the 20 minutes before optogenetic perturbation.

### FAK lifetime analysis

To analyze focal adhesion dynamics, individual focal adhesions were manually identified and labeled. To determine their lifetime, only those focal adhesions that were both formed and dissolved during the 1h timelapse recording were considered.

### Tracking of cell nuclei

Nuclear movements were measured via the automated tracking tool ‘TrackMate’, which is available for Fiji.^51,52^ For 2D tracking, cells were stained with Spy650 Live-cell DNA stain (1:1000 dilution, 1.5 h) (Spirochrome) before microscopy. To track nuclear movements in 3D spheroids, opto-control and opto-GEF-H1 cells that stably express a labeled Histone (mCherry-H1) were used for imaging. First, the nuclei were detected using the implemented LoG detector with a quality threshold (2D: 0.651; 3D: 3) and an estimated object diameter (2D: 15 μm; 3D: 12 μm). The detected nuclei were then filtered based on their quality and tracks were generated using the simple LAP tracker with a specified linking max distance (2D: 5 μm; 3D: 2 μm), Gap-closing max distance (2D: 15 μm; 3D: 2 μm) and Gap-closing max frame gap of 2. The resulting tracks were then filtered again to exclude incomplete tracks that covered less than 99.5% (2D) or 85% (3D) of the time-lapse movie, respectively. For 3D spheroids, the average movement velocities of cells within one spheroid were averaged for each time point.

### Semi-automated spheroid area analysis

To evaluate changes in spheroid size, the projected 2D spheroid area was measured. First, the spheroid was identified in each fluorescence image of a timelapse series via Otsu thresholding. The thresholded area was measured and normalized to the average area of all timepoints before optogenetic perturbation. In brightfield timelapse series the spheroid area was tracked manually before and at the end of the optogenetic perturbation.

### Statistical analysis

All plots and statistical analyses were performed using Prism (GraphPad). Image panels were created using ImageJ/Fiji.^50^ The type of statistical tests and significance levels are indicated in the respective figure legends. P-values are indicated by stars (*: P<0.05; **: P<0.01; ***: P<0.001; ****: P<0.0001).

## Acknowledgments

We acknowledge the use of the imaging equipment and the support in microscope usage by the “Imaging Center Campus Essen” (ICCE), Center of Medical Biotechnology (ZMB), University of Duisburg-Essen. This work was funded by the Deutsche Forschungsgemeinschaft (DFG, German Research Foundation) Priority Programs SPP 1926 under the grant number NA 413/4-1 to P.N., L.D. and J.W., SFB 1430 – Project-ID 424228829 to P.N., N.S., N.Sc. and A.P., the DFG Heisenberg Program grants DE 823/6-1, 823/8-1 to L.D. and the Principal Investigator grant DE 823/10-1 to L.D. The Nikon Ti2-E TIRF DualCam microscope was funded by the Deutsche Forschungsgemeinschaft (DFG, German Research Foundation) – Project-ID 361032973.

## Online supplemental material

### Supplementary Figures

**Figure S1:**
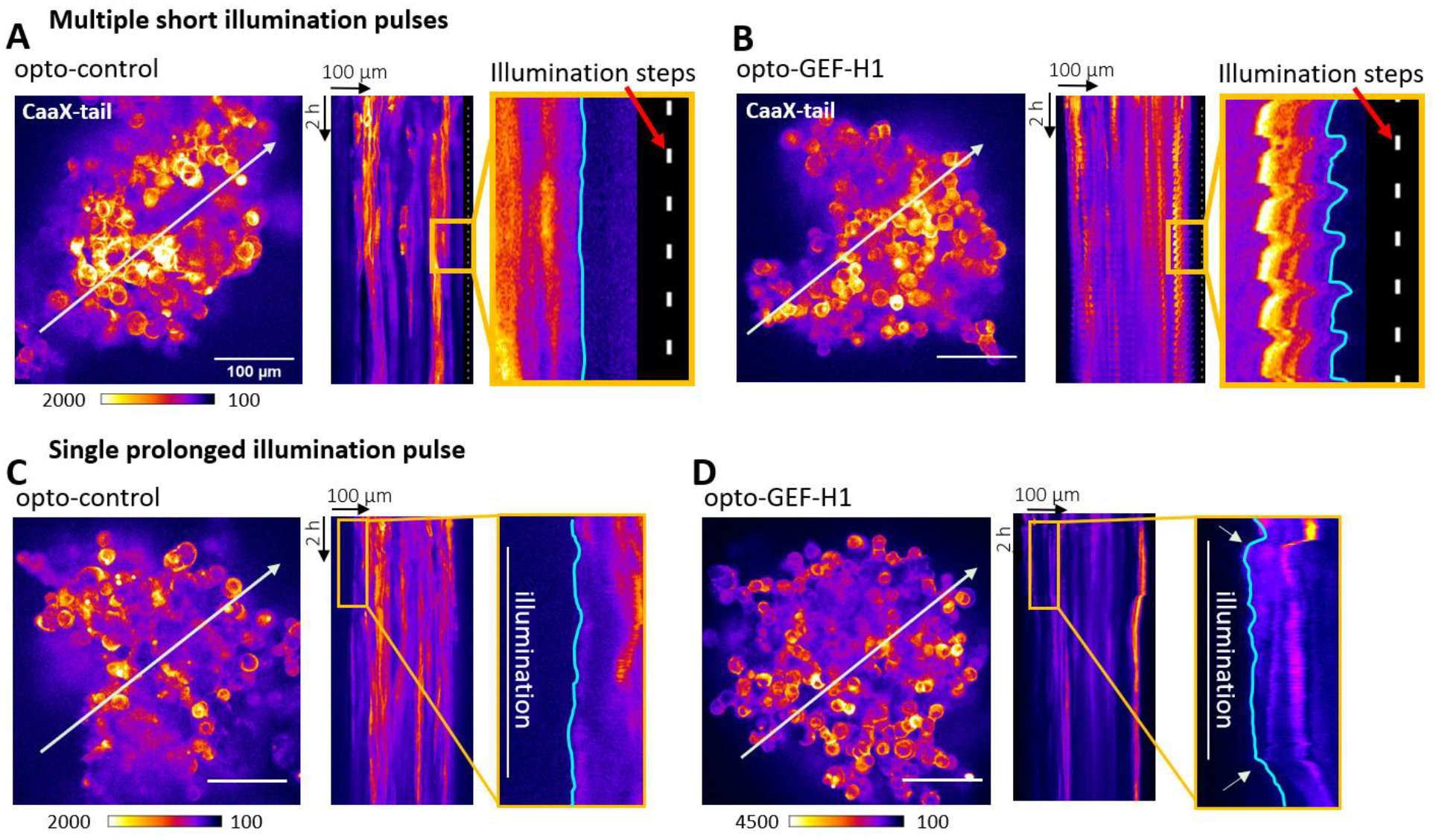
Multicellular melanoma spheroids expand in a pulsatory manner upon externally controlled, synchronous contraction dynamics, and stably upon stimulation of self-organized asynchronous contraction dynamics. (A-D) Left panels: Representative spinning disk microscopy images of the CaaX-tail plasma membrane marker in B16F1-opto-control (A, C) and B16F1-opto-GEF-H1 (B, D) spheroids. Right panels: Corresponding kymograph analysis along the white arrows in left panels. The zoomed in regions are indicated by orange boxes. Optogenetic stimulations are indicated by white lines in the zoomed kymograph regions. A representative spheroid is shown for each condition, taken from 3 opto-GEF-H1 and 3 opto-control spheroids examined in 3 independent experiments. Spheroids were imaged over a total time period of 12.5 h while applying 36 individual 5 min lasting optogenetic stimulation pulses with 15 min intermittent pauses (A, B) or over a total time period of 12.5 h while applying a single, 3 h lasting optogenetic stimulation pulse followed by 9 h of recovery (C, D).

**Figure S2:**
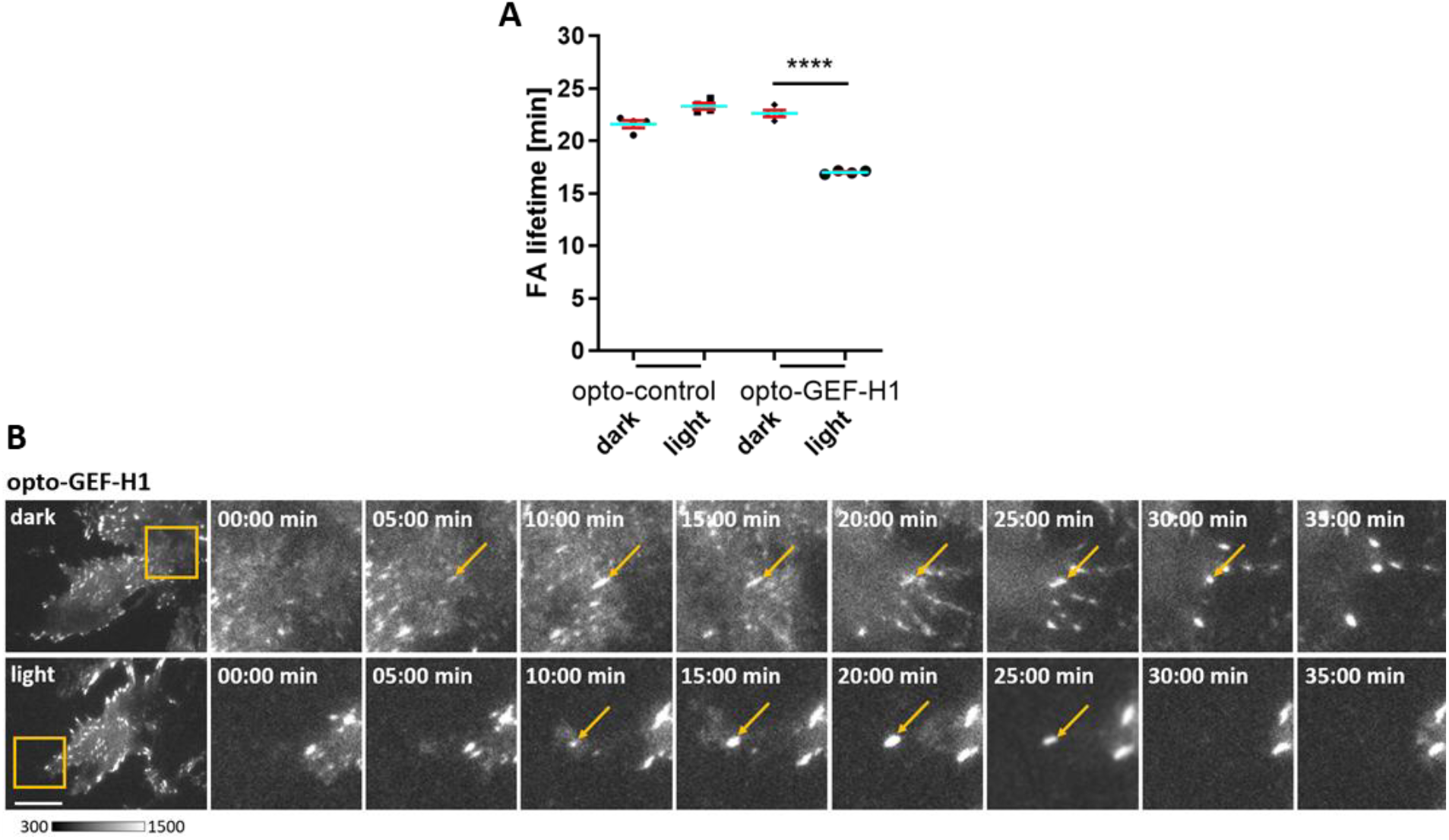
Optogenetic stimulation of cell contraction dynamics increases focal adhesion turn-over. (A) Quantification of average lifetime of dynamic focal adhesions (FAs) in adherent B16F1-opto-control and B16F1-opto-GEF-H1 with and without optogenetic stimulation. B16F1-opto-control N=142 FAs (dark; 44 cells), N=144 FAs (light; 44 cells), B16F1-opto-GEF-H1 N= 123 FAs (dark; 24 cells), N=122 FAs (light; 26 cells) from 4 independent experiments. Paired t-test. Error bars represent S.E.M. (B) Representative TIRF images of B16F1-opto-GEF-H1 cells expressing paxillin (pmCherry-Paxillin mouse) during the prerun phase (upper row) and during optogenetic stimulation (light, lower row). Scale bar=20 μm.

**Figure S3:**
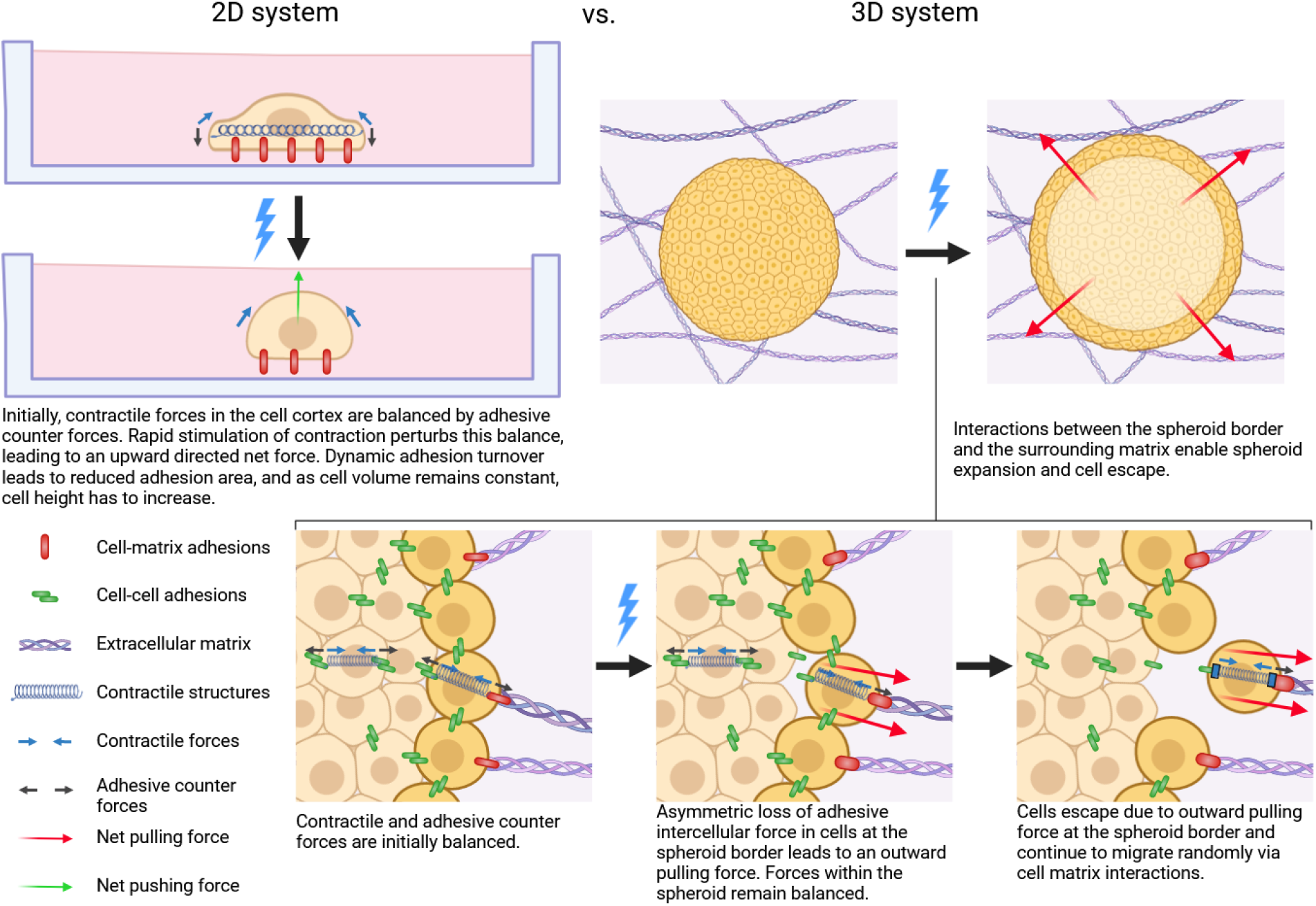
Morphological responses to optogenetically stimulated cell contraction dynamics. Top panels: Cells that adhere to a surface in 2D cell culture reduce their adhesive cell area upon optogenetic stimulation of contraction dynamics, leading to a rounded morphology. Multicellular melanoma spheroids (3D cell culture) expand upon optogenetic stimulation of contraction dynamics. Bottom panels show a proposed mechanism that could explain the experimental observations.

### Supplementary Videos

Video 1: **Enhanced cell contraction dynamics in B16F1 mouse melanoma cells upon increased expression of a constitutively active GEF-H1 mutant**. TIRF videos of representative cells transiently expressing myosin IIa (pCMV-mCherry-MHC IIA) and the constitutively active GEF-H1 C53R mutant (pCMV5-EGFP-GEF-HI C53R) or a control vector (pEGFP-N1), respectively. Frame rate: 3/min. Scale bar=20 μm.

Video 2: **Optogenetic stimulation of a single cell contraction pulse in adherent mouse melanoma cells using a single, short illumination pulse**. TIRF videos of myosin IIa signal and opto-control (left) or opto-GEF-H1 (right) with 5 min optogenetic stimulation at 445-488 nm. Frame rate: 3/min. Scale bar=20 μm.

Video 3: **Optogenetic stimulation of multiple autonomous cell contraction pulses using a longer, single illumination pulse**. TIRF videos of myosin IIa signal and opto-control (left) or opto-GEF-H1 (right) with 20 min optogenetic stimulation at 445-488 nm. Frame rate: 3/min. Scale bar=20 μm.

Video 4: **Enhanced cell movement dynamics within opto-GEF-H1 spheroids during optogenetic stimulation**. Spinning disk confocal microscopy videos of nuclei (Histone-H1) in B16F1-opto-control or B16F1-opto-GEF-H1 spheroids with a short (5 min) optogenetic stimulation. Frame rate: 3/min. Scale bar=100 μm.

Video 5: **Increase of opto-GEF-H1 spheroid area upon optogenetic GEF-H1 release**. Epifluorescence videos of nuclei (Histone-H1) in B16F1-opto-control or B16F1-opto-GEF-H1 spheroids with a short (5 min) optogenetic stimulation. Frame rate: 3/min. Scale bar=100 μm.

Video 6: **Escape of individual cells from melanoma spheroids upon long-term stimulation of cell contraction dynamics**. Bright field microscopy movies of opto-control and opto-GEF-H1 spheroids embedded in collagen matrix during continuous optogenetic stimulation for 12 h. Frame rate: 1/min. Scale bar=100 μm.

## Notes

### Competing Interest Statement

The authors have declared no competing interest.

## References

1. Lauffenburger, D.A., and Horwitz, A.F. (1996). Cell migration: a physically integrated molecular process. Cell 84, 359–369.

2. Yamaguchi, H., and Condeelis, J. (2007). Regulation of the actin cytoskeleton in cancer cell migration and invasion. Biochimica et Biophysica Acta (BBA) - Molecular Cell Research 1773, 642–652. 10.1016/j.bbamcr.2006.07.001.

3. Olson, M.F., and Sahai, E. (2009). The actin cytoskeleton in cancer cell motility. Clin Exp Metastasis 26, 273. 10.1007/s10585-008-9174-2.

4. Fackler, O.T., and Grosse, R. (2008). Cell motility through plasma membrane blebbing. J Cell Biol 181, 879–884. 10.1083/jcb.200802081.

5. Itoh, K., Yoshioka, K., Akedo, H., Uehata, M., Ishizaki, T., and Narumiya, S. (1999). An essential part for Rho–associated kinase in the transcellular invasion of tumor cells. Nat Med 5, 221–225. 10.1038/5587.

6. Kaneko, K., Satoh, K., Masamune, A., Satoh, A., and Shimosegawa, T. (2002). Myosin Light Chain Kinase Inhibitors Can Block Invasion and Adhesion of Human Pancreatic Cancer Cell Lines. Pancreas 24, 34–41. 10.1097/00006676-200201000-00005.

7. Liu, S., Goldstein, R.H., Scepansky, E.M., and Rosenblatt, M. (2009). Inhibition of Rho-Associated Kinase Signaling Prevents Breast Cancer Metastasis to Human Bone. Cancer Res 69, 8742–8751. 10.1158/0008-5472.CAN-09-1541.

8. Duxbury, M.S., Ashley, S.W., and Whang, E.E. (2004). Inhibition of pancreatic adenocarcinoma cellular invasiveness by blebbistatin: a novel myosin II inhibitor. Biochem Biophys Res Commun 313, 992–997. 10.1016/j.bbrc.2003.12.031.

9. Ivkovic, S., Beadle, C., Noticewala, S., Massey, S.C., Swanson, K.R., Toro, L.N., Bresnick, A.R., Canoll, P., and Rosenfeld, S.S. (2012). Direct inhibition of myosin II effectively blocks glioma invasion in the presence of multiple motogens. Mol Biol Cell 23, 533–542. 10.1091/mbc.e11-01-0039.

10. Graessl, M., Koch, J., Calderon, A., Kamps, D., Banerjee, S., Mazel, T., Schulze, N., Jungkurth, J.K., Patwardhan, R., Solouk, D., et al. (2017). An excitable Rho GTPase signaling network generates dynamic subcellular contraction patterns. J Cell Biol 216, 4271–4285. 10.1083/jcb.201706052.

11. Kamps, D., Koch, J., Juma, V.O., Campillo-Funollet, E., Graessl, M., Banerjee, S., Mazel, T., Chen, X., Wu, Y.-W., Portet, S., et al. (2020). Optogenetic Tuning Reveals Rho Amplification-Dependent Dynamics of a Cell Contraction Signal Network. Cell Rep 33, 108467. 10.1016/j.celrep.2020.108467.

12. Cheng, I.K., Tsang, B.C., Lai, K.P., Ching, A.K., Chan, A.W., To, K., Lai, P.B., and Wong, N. (2012). GEF-H1 over-expression in hepatocellular carcinoma promotes cell motility via activation of RhoA signalling. J Pathol 228, 575–585. 10.1002/path.4084.

13. Liao, Y.C., Ruan, J.W., Lua, I., Li, M.H., Chen, W.L., Wang, J.R., Kao, R.H., and Chen, J.H. (2012). Overexpressed hPTTG1 promotes breast cancer cell invasion and metastasis by regulating GEF-H1/RhoA signalling. Oncogene 31, 3086–3097. 10.1038/onc.2011.476.

14. Tao Kai, Guo, B., Zhang, Y., Hui, Q., Chang, P., and Shi, J. (2016). Guanine nucleotide exchange factor H1 can be a new biomarker of melanoma. Biologics Volume 10, 89–98. 10.2147/BTT.S109643.

15. Joo, E., and Olson, M.F. (2021). Regulation and functions of the RhoA regulatory guanine nucleotide exchange factor GEF-H1. Small GTPases 12, 358–371. 10.1080/21541248.2020.1840889.

16. Cao, J., Yang, T., Tang, D., Zhou, F., Qian, Y., and Zou, X. (2019). Increased expression of GEF-H1 promotes colon cancer progression by RhoA signaling. Pathol Res Pract 215, 1012–1019. 10.1016/j.prp.2019.02.008.

17. Birkenfeld, J., Nalbant, P., Yoon, S.-H., and Bokoch, G.M. (2008). Cellular functions of GEF-H1, a microtubule-regulated Rho-GEF: is altered GEF-H1 activity a crucial determinant of disease pathogenesis? Trends Cell Biol 18, 210–219. 10.1016/j.tcb.2008.02.006.

18. Ding, Z., Dong, Z., Yang, Y., Fortin Ensign, S.P., Sabit, H., Nakada, M., Ruggieri, R., Kloss, J.M., Symons, M., Tran, N.L., et al. (2020). Leukemia-Associated Rho Guanine Nucleotide Exchange Factor and Ras Homolog Family Member C Play a Role in Glioblastoma Cell Invasion and Resistance. Am J Pathol 190, 2165–2176. 10.1016/j.ajpath.2020.07.005.

19. Yu, H.-G., Nam, J.-O., Miller, N.L.G., Tanjoni, I., Walsh, C., Shi, L., Kim, L., Chen, X.L., Tomar, A., Lim, S.-T., et al. (2011). p190RhoGEF (Rgnef) Promotes Colon Carcinoma Tumor Progression via Interaction with Focal Adhesion Kinase. Cancer Res 71, 360–370. 10.1158/0008-5472.CAN-10-894.

20. Cervantes-Villagrana, R.D., García-Jiménez, I., and Vázquez-Prado, J. (2023). Guanine nucleotide exchange factors for Rho GTPases (RhoGEFs) as oncogenic effectors and strategic therapeutic targets in metastatic cancer. Cell Signal 109, 110749. 10.1016/j.cellsig.2023.110749.

21. Ren, Y., Li, R., Zheng, Y., and Busch, H. (1998). Cloning and Characterization of GEF-H1, a Microtubule-associated Guanine Nucleotide Exchange Factor for Rac and Rho GTPases. Journal of Biological Chemistry 273, 34954–34960. 10.1074/jbc.273.52.34954.

22. Krendel, M., Zenke, F.T., and Bokoch, G.M. (2002). Nucleotide exchange factor GEF-H1 mediates cross-talk between microtubules and the actin cytoskeleton. Nat Cell Biol 4, 294–301. 10.1038/ncb773.

23. Chang, Y.-C., Nalbant, P., Birkenfeld, J., Chang, Z.-F., and Bokoch, G.M. (2008). GEF-H1 Couples Nocodazole-induced Microtubule Disassembly to Cell Contractility via RhoA. Mol Biol Cell 19, 2147–2153. 10.1091/mbc.e07-12-1269.

24. Wang, H., and Hahn, K.M. (2016). LOVTRAP: A Versatile Method to Control Protein Function with Light. Curr Protoc Cell Biol 73. 10.1002/cpcb.12.

25. Amano, M., Ito, M., Kimura, K., Fukata, Y., Chihara, K., Nakano, T., Matsuura, Y., and Kaibuchi, K. (1996). Phosphorylation and activation of myosin by Rho-associated kinase (Rho-kinase). Journal of Biological Chemistry 271, 20246–20249. 10.1074/jbc.271.34.20246.

26. Kimura, K., Ito, M., Amano, M., Chihara, K., Fukata, Y., Nakafuku, M., Yamamori, B., Feng, J., Nakano, T., Okawa, K., et al. (1996). Regulation of Myosin Phosphatase by Rho and Rho-Associated Kinase (Rho-Kinase). Science (1979) 273, 245–248. 10.1126/SCIENCE.273.5272.245.

27. Nalbant, P., Wagner, J., and Dehmelt, L. (2023). Direct investigation of cell contraction signal networks by light-based perturbation methods. Pflugers Arch 475, 1439–1452. 10.1007/s00424-023-02864-2.

28. Kamps, D., Koch, J., Juma, V.O., Campillo-Funollet, E., Graessl, M., Banerjee, S., Mazel, T., Chen, X., Wu, Y.W., Portet, S., et al. (2020). Optogenetic Tuning Reveals Rho Amplification-Dependent Dynamics of a Cell Contraction Signal Network. Cell Rep 33, 108467. 10.1016/j.celrep.2020.108467.

29. Smalley, K.S., Lioni, M., Noma, K., Haass, N.K., and Herlyn, M. (2008). In vitro three-dimensional tumor microenvironment models for anticancer drug discovery. Expert Opin Drug Discov 3, 1–10. 10.1517/17460441.3.1.1.

30. Mierke, C.T., Rösel, D., Fabry, B., and Brábek, J. (2008). Contractile forces in tumor cell migration. Eur J Cell Biol 87, 669–676. 10.1016/j.ejcb.2008.01.002.

31. Provenzano, P.P., and Keely, P.J. (2009). The role of focal adhesion kinase in tumor initiation and progression. Cell Adh Migr 3, 347–350. 10.4161/cam.3.4.9458.

32. Schlaepfer, D.D., Mitra, S.K., and Ilic, D. (2004). Control of motile and invasive cell phenotypes by focal adhesion kinase. Biochimica et Biophysica Acta (BBA) - Molecular Cell Research 1692, 77–102. 10.1016/j.bbamcr.2004.04.008.

33. Billion, K. (2006). Proteolytic cleavage of cadherins: Functional role of the cleaved extracellular and cytoplasmic domains.

34. Plotnikov, S. V, and Waterman, C.M. (2013). Guiding cell migration by tugging. Curr Opin Cell Biol 25, 619–626. 10.1016/j.ceb.2013.06.003.

35. Nalbant, P., and Dehmelt, L. (2018). Exploratory cell dynamics: a sense of touch for cells? Biol Chem 399, 809–819. 10.1515/hsz-2017-0341.

36. Plotnikov, S. V, Pasapera, A.M., Sabass, B., and Waterman, C.M. (2012). Force fluctuations within focal adhesions mediate ECM-rigidity sensing to guide directed cell migration. Cell 151, 1513–1527. 10.1016/j.cell.2012.11.034.

37. Reuther, G.W., Lambert, Q.T., Booden, M.A., Wennerberg, K., Becknell, B., Marcucci, G., Sondek, J., Caligiuri, M.A., and Der, C.J. (2001). Leukemia-associated Rho Guanine Nucleotide Exchange Factor, a Dbl Family Protein Found Mutated in Leukemia, Causes Transformation by Activation of RhoA. Journal of Biological Chemistry 276, 27145–27151. 10.1074/jbc.M103565200.

38. Yan, S., Xue, H., Zhang, P., Han, X., Guo, X., Yuan, G., Deng, L., and Li, G. (2016). MMP inhibitor Ilomastat induced amoeboid-like motility via activation of the Rho signaling pathway in glioblastoma cells. Tumor Biology 37, 16177–16186. 10.1007/s13277-016-5464-5.

39. Friedl, P., Borgmann, S., and Bröcker, E.-B. (2001). Amoeboid leukocyte crawling through extracellular matrix: lessons from the Dictyostelium paradigm of cell movement. J Leukoc Biol, 70(4):491–509.

40. Eitaki, M., Yamamori, T., Meike, S., Yasui, H., and Inanami, O. (2012). Vincristine enhances amoeboid-like motility via GEF-H1/RhoA/ROCK/Myosin light chain signaling in MKN45 cells. BMC Cancer 12, 469. 10.1186/1471-2407-12-469.

41. Liu, Y.-J., Le Berre, M., Lautenschlaeger, F., Maiuri, P., Callan-Jones, A., Heuzé, M., Takaki, T., Voituriez, R., and Piel, M. (2015). Confinement and Low Adhesion Induce Fast Amoeboid Migration of Slow Mesenchymal Cells. Cell 160, 659–672. 10.1016/j.cell.2015.01.007.

42. Blaser, H., Reichman-Fried, M., Castanon, I., Dumstrei, K., Marlow, F.L., Kawakami, K., Solnica-Krezel, L., Heisenberg, C.-P., and Raz, E. (2006). Migration of Zebrafish Primordial Germ Cells: A Role for Myosin Contraction and Cytoplasmic Flow. Dev Cell 11, 613–627. 10.1016/j.devcel.2006.09.023.

43. Lämmermann, T., and Sixt, M. (2009). Mechanical modes of ‘amoeboid’ cell migration. Curr Opin Cell Biol 21, 636–644. 10.1016/j.ceb.2009.05.003.

44. Krendel, M., Zenke, F.T., and Bokoch, G.M. (2002). Nucleotide exchange factor GEF-H1 mediates cross-talk between microtubules and the actin cytoskeleton. Nat Cell Biol 4, 294–301. 10.1038/ncb773.

45. Dulyaninova, N.G., House, R.P., Betapudi, V., and Bresnick, A.R. (2007). Myosin-IIA Heavy-Chain Phosphorylation Regulates the Motility of MDA-MB-231 Carcinoma Cells. Mol Biol Cell 18, 3144–3155. 10.1091/mbc.e06-11-1056.

46. Wang, W., Lin, C., Lu, D., Ning, Z., Cox, T., Melvin, D., Wang, X., Bradley, A., and Liu, P. (2008). Chromosomal transposition of PiggyBac in mouse embryonic stem cells. Proceedings of the National Academy of Sciences 105, 9290–9295. 10.1073/pnas.0801017105.

47. Costantini, L.M., Baloban, M., Markwardt, M.L., Rizzo, M.A., Guo, F., Verkhusha, V. V., and Snapp, E.L. (2015). A palette of fluorescent proteins optimized for diverse cellular environments. Nat Commun 6, 7670. 10.1038/ncomms8670.

48. Adachi, K., Kopp, W., Wu, G., Heising, S., Greber, B., Stehling, M., Araúzo-Bravo, M.J., Boerno, S.T., Timmermann, B., Vingron, M., et al. (2018). Esrrb Unlocks Silenced Enhancers for Reprogramming to Naive Pluripotency. Cell Stem Cell 23, 266–275.e6. 10.1016/j.stem.2018.05.020.

49. Morgenstern, J.P., and Land, H. (1990). Advanced mammalian gene transfer: high titre retroviral vectors with multiple drug selection markers and a complementary helper-free packaging cell line. Nucleic Acids Res 18, 3587–3596. 10.1093/nar/18.12.3587.

50. Schindelin, J., Arganda-Carreras, I., Frise, E., Kaynig, V., Longair, M., Pietzsch, T., Preibisch, S., Rueden, C., Saalfeld, S., Schmid, B., et al. (2012). Fiji: an open-source platform for biological-image analysis. Nat Methods 9, 676–682. 10.1038/nmeth.2019.

51. Ershov, D., Phan, M.-S., Pylvänäinen, J.W., Rigaud, S.U., Le Blanc, L., Charles-Orszag, A., Conway, J.R.W., Laine, R.F., Roy, N.H., Bonazzi, D., et al. (2022). TrackMate 7: integrating state-of-the-art segmentation algorithms into tracking pipelines. Nat Methods 19, 829–832. 10.1038/s41592-022-01507-1.

52. Tinevez, J.Y., Perry, N., Schindelin, J., Hoopes, G.M., Reynolds, G.D., Laplantine, E., Bednarek, S.Y., Shorte, S.L., and Eliceiri, K.W. (2017). TrackMate: An open and extensible platform for single-particle tracking. Methods 115, 80–90. 10.1016/J.YMETH.2016.09.016.

